# Generation of ENSEMBL-based proteogenomics databases boosts the identification of non-canonical peptides

**DOI:** 10.1101/2021.06.08.447496

**Authors:** Husen M. Umer, Yafeng Zhu, Julianus Pfeuffer, Timo Sachsenberg, Janne Lehtiö, Rui Branca, Yasset Perez-Riverol

## Abstract

**Summary:** We have implemented the *pypgatk* package and the *pgdb* workflow to create proteogenomics databases based on ENSEMBL resources. The tools allow the generation of protein sequences from novel protein-coding transcripts by performing a three-frame translation of pseudogenes, lncRNAs, and other non-canonical transcripts, such as those produced by alternative splicing events. It also includes exonic out-of-frame translation from otherwise canonical protein-coding mRNAs. Moreover, the tool enables the generation of variant protein sequences from multiple sources of genomic variants including COSMIC, cBioportal, gnomAD, and mutations detected from sequencing of patient samples. *pypgatk* and *pgdb* provide multiple functionalities for database handling, notably optimized target/decoy generation by the algorithm *DecoyPyrat*. Finally, we perform a reanalysis of four public datasets in PRIDE by generating cell-type specific databases for 65 cell lines using the *pypgatk* and *pgdb* workflow, revealing a wealth of non-canonical or cryptic peptides amounting to more than 10% of the total number of peptides identified (43,501 out of 402,512).

**Availability:** The software is freely available. *pypgatk*: (https://github.com/bigbio/py-pgatk/), and *pgdb*: (https://github.com/nf-core/pgdb)

**Contact:** Yasset Perez-Riverol, Rui Branca

**Supplementary information:** Supplementary data are available online.

## 1 Introduction

Proteogenomics is a rapidly developing multi-omics field that integrates genomics and transcriptomics information with shotgun Proteomics Mass Spectrometry data to improve gene annotation, often uncovering novel or non-canonical protein coding regions in the genome (Branca, *et al.*, 2014). One of the most important applications is in the study of cancer cells and tumors, where identifying aberrant proteins holds great potential in both elucidating cancer biology and in developing cancer therapies. However, the discovery of these aberrant proteins remains particularly challenging and is still largely linked to evidence from the sequencing level, rather than directly from the protein data level. Proteomics of large datasets has exploded in recent years with many large datasets now available in public repositories. Recent applications of proteogenomics have enabled multiomics detection of novel peptide sequences that are not present in the canonical protein database. For example, Cuevas et al recently identified a large number of non-canonical proteins in B cell lymphomas (Ruiz Cuevas, *et al.*, 2021). To solve the bottleneck of performing high throughput proteogenomics analysis, we present a *Python* application integrated into a Nextflow workflow for large-scale generation of proteogenomics databases enabling the identification and quantification of variant proteins (derived from single nucleotide variant mutations) and non-canonical or cryptic proteins (from normally dormant regions of the genome).

## 2 Implementation

We implemented *pypgatk*, a *Python* package that provides tools to generate protein databases from non-canonical sequences as well as DNA variants and mutations from public resources as well as custom files (**Figure 1a**).

**Figure 1:**
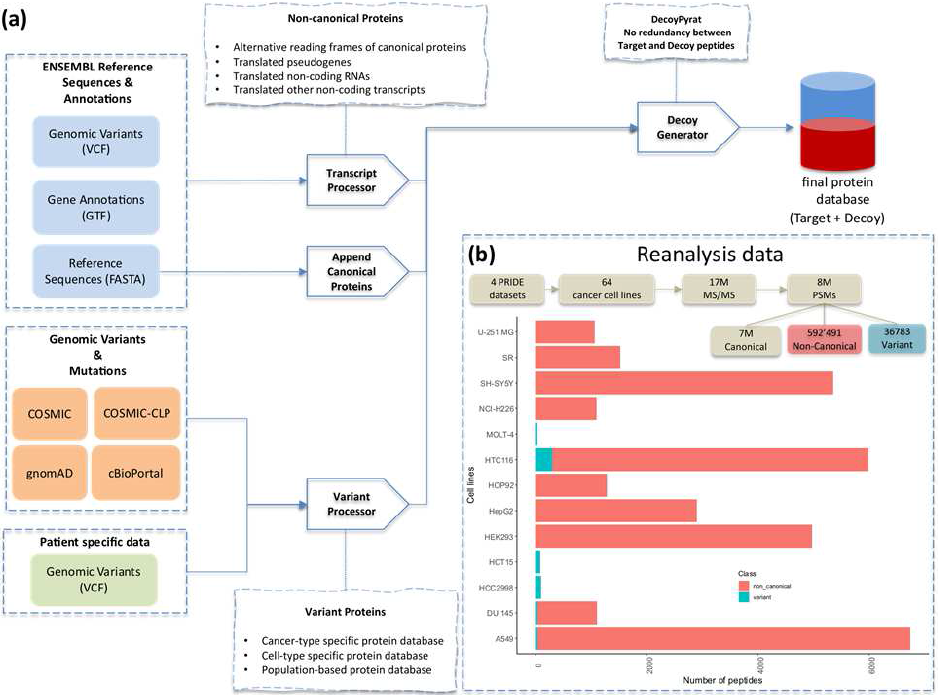
**(a)** *pypgatk* and *pgdb* components to generate ENSEMBL-based proteogenomics databases. **(b)** Distribution of novel and variant peptides identified using proteogeonomics databases and four cancer cell lines’ datasets from PRIDE.

### Non-canonical protein databases

Non-canonical proteins are a product of translation of transcripts that are not reported as protein-coding in the reference protein databases, or a product of out-of-frame translation of canonical transcripts. While many of the non-canonical proteins could be attributed to the yet incomplete reference databases, they might also be attributed to activation of those genes under certain conditions such as genetic and epigenetic misregulation in cancer (Zhu, **et al.**, 2018). We have developed the ***dnaseq-to-proteindb*** tool to generate protein sequences from non-canonical protein-coding transcripts by performing a three-frame translation of pseudogenes, lncRNAs, etc. It also extracts alternative reading frames from canonical protein-coding genes to enable the detection of out-of-frame cryptic proteins (Vanderperre, *et al.*, 2013). Furthermore, the ***ensemble-downloader*** tool enables automatic download of the latest ENSEMBL (Yates, **et al.**, 2020) resources including gene annotations, the reference genome, and canonical proteins for the species of interest.

### Variant protein databases

Detection of altered proteins from proteomics data requires the inclusion of the mutated sequences in the target databases. However, due to a large number of potential DNA variants, only potentially relevant variant sequences should be included to keep the database size under control. Here, we implemented methods to enable the automatic generation of variant proteins from publicly available cancer mutations datasets, cancer cell lines, and custom VCF files obtained from genome sequencing. ***cosmic-to-proteindb*** and ***cbioportal-to-proteindb*** enable the generation of cancer-type specific protein databases by generating mutated protein sequences absed on genomic mutations identified in cancer samples. ***cosmic-to-proteindb*** curates mutations from the Catalogue Of Somatic Mutations In Cancer (COSMIC). It allows filtering the mutations based on cancer type or tissue of origin. Alternatively, ***cbioportal-to-proteindb*** translates genomics mutations reported by thousands of cancer studies through cBioPortal. *pypgatk* enables downloading and processing mutations from ENSEMBL and gnomAD resources. ***vcf-to-proteindb*** translates the genomic variants into variant protein sequences. The variants can be filtered based on minor allele frequency to enable a special focus on common variants. ***vcf-to-proteindb*** command accepts a custom VCF file from sample-specific datasets and generates a database of altered protein-coding sequences, allowing the generation of proteogenomics databases from the samples when whole exome or whole genome sequencing data is available.

## 3 Large scale analysis

To enable large-scale processing of genomics data to generate protein databases, we have also built the Proteomics-Genomics DataBase (***pgdb*** - https://nf-co.re/pgdb) workflow in *Nextflow using bioconda and BioContainers*. The pipeline integrates the various commands of *pypgatk* allowing the user to generate protein databases by simple parameter selection without any additional input required from the user.

### Identification of non-canonical peptides

We applied ***pgdb*** to generate cell-type specific databases for 63 cell lines (**Supplementary Note 1**). Mutations from the COSMIC Cell Line project and the Broad CCLE project through cBioPortal were downloaded for each cell line to generate the respective set of variant protein sequences. Additionally, a database of non-canonical proteins was generated from the latest human genome assembly. The variant protein database from each cell line was appended to the non-canonical and canonical protein databases and the decoy sequences were generated to search MS/MS proteomics datasets from the corresponding cell lines. The proteomics data were obtained through PRIDE database (Perez-Riverol, *et al.*, 2019) (PXD005946, PXD019263, PXD004452, and PXD014145). proteomicsLFQ (https://nf-co.re/proteomicslfq) was used to identify the novel peptides (Supplementary Note 1-4). Overall, 402,512 target peptide sequences were identified, including 43,501 non-canonical peptides and 786 variant peptide sequences (**Figure 1b**). The majority of non-canonical peptides were novel coding sequences whereas only 16% were matching canonical protein sequences with at least one amino acid mismatch.

## 4 Conclusions

The developed tools facilitate the creation of proteogenomics databases based on ENSEMBL genomes and other relevant sources of genome variation information. The *pgdb* is the first workflow for proteogenomics database generation and its development within the nf-core community will ensure its stability, continued development, and community support. ***pyp-gatk*** (https://pgatk.readthedocs.io/en/latest/pypgatk.html) and ***pgdb*** (https://nf-co.re/pgdb/1.0.0/usage) include extensive documentation to help bioinformaticians and biologists to create their custom proteogenomics databases.

