## Supplementary Notes for "Generation of ENSEMBL-based proteogenomics databases boosts the identification of non-canonical peptides"

[1] Department of Oncology Pathology, Science for Life Laboratory, Karolinska Institutet, Solna, Stockholm 17121, Sweden. [2] Department of Genetics, Harvard Medical School, Boston, MA 02115 USA. [3] Algorithmic Bioinformatics, Freie Universität Berlin, Berlin, Germany; Visualization and Data analysis, Zuse Institute Berlin, Berlin, Germany. [4] Institute for Bioinformatics and Medical Informatics, University of Tübingen, Sand 14, 72076, Tübingen, Germany. [5] European Molecular Biology Laboratory, European Bioinformatics Institute (EMBL-EBI), Wellcome Trust Genome Campus, Hinxton, Cambridge, CB10 1SD, UK.

**Supplementary Note 1**: Reanalysed datasets description

| PX accession | # Samples reanalyzed | Cell lines | # MS/MS | PSMs | Quantified Peptides |
|---|---|---|---|---|---|
| PXD004452 (Bekker-Jensen, et al., 2017) | 4 | HTC116, HEK293, SH-SY5Y, A549 | 4’028’557 | 1’744’509 | 1’474’088 |
| PXD005946 (Gholami, et al., 2013) | 61 | NCI-H522, HCC2998, SN12C, HS578T, K562, BT549, EKVX, HOP62, SK-MEL-28, T47D, MDAMB435, SR, MDAMB231, LOXIMVI, SNB-75, M14, MCF7/AdrR, HT-29, NCI-H226, RXF 393 U-251 MG, SK-MEL-5 SKOV3, DU 145 OVCAR3, IGROV-1 SK-MEL-2, OVCAR4 UACC-62, COLO205 UACC-257, SNB-19 786-0, HL-60 SW620, A549, A498 CCRFCEM, NCI-H23 Malme3M, UO31 TK-10, HOP92 HCT116, NCI-H322M ACHN, HCT15, Caki1 SF539, RPMI8226, SF295, MOLT-4, MCF-7 OVCAR5, SF268 NCI-H460, OVCAR8, PC-3, KM12 | 11’765’655 | 6’063’277 | 2’870’404 |
| PXD014145 (Baumert, et al., 2020) | 1 | A549 | 331’365 | 138’639 | 131’078 |
| PXD019263 (Vavilov, et al., 2020) | 2 | HepG2 | 896’891 | 502’102 | 156’289 |
| Total | 68 | 63 cell lines | 17’022’468 | 8’448’527 | 4’631’859 |

**Supplementary Note 2:** Distribution of search engine score vs peptide class

The distribution of search engine score vs the peptide class: ENSP (ENSEMBL proteins), OTHER-REFS (peptides mapping to Uniprot, RefSeq, or GENCODE), contaminant, non-canonical, and variants. The distribution of scores for variant peptides is similar to canonical peptides. However, the number of non-canonical peptides with score lower than 0.99 is higher than for other classes of peptides.

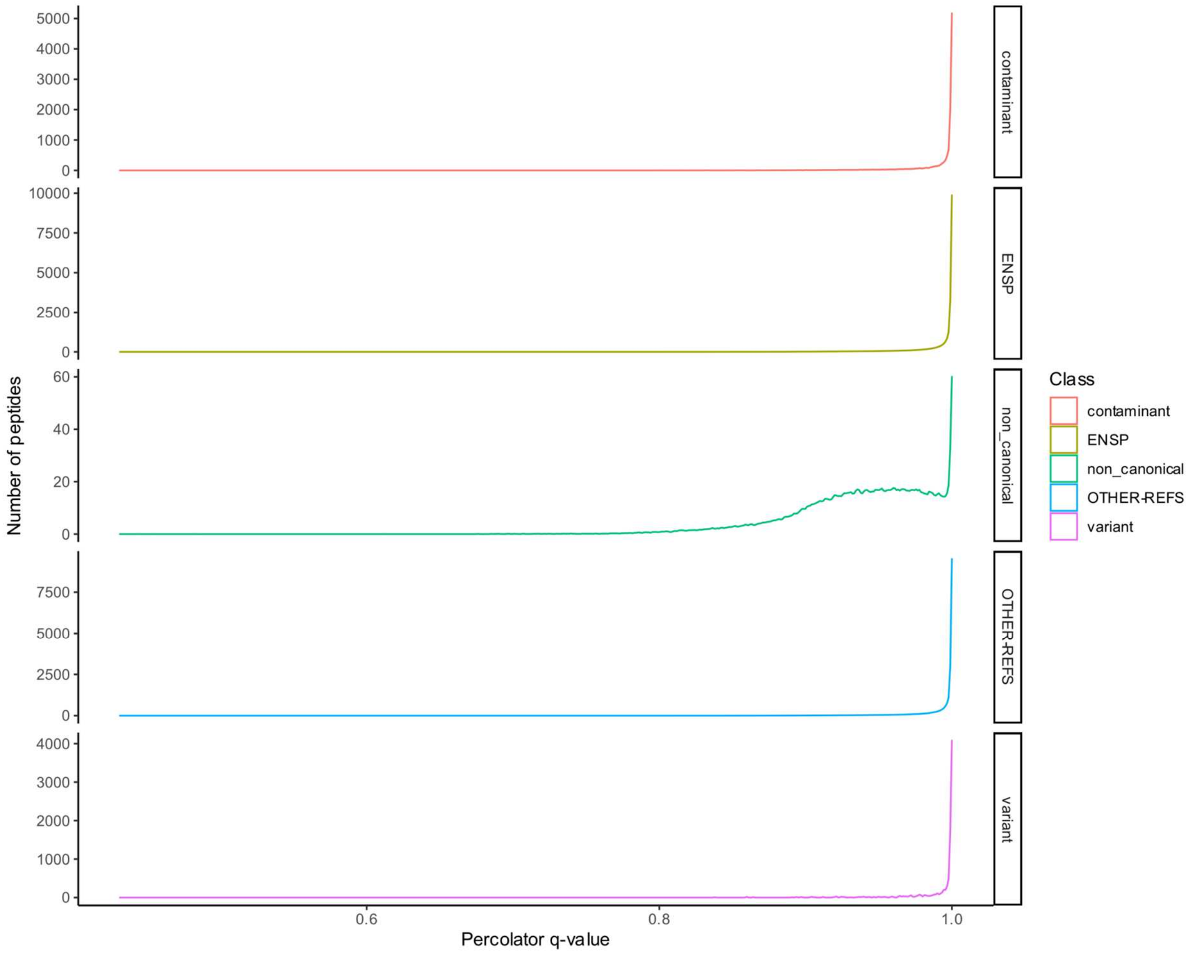

**Supplementary Note 3**: Distribution of number amino acids per peptide sequence

Distribution of number of amino acids (aa) per peptide sequence for the for different classes of peptides: ENSP (ENSEMBL proteins), OTHER-REFS (peptides mapping to Uniprot, RefSeq, or GENCODE), contaminant, non-canonical, and variants. On average, canonical peptides (ENSP and OTHER-REFS) peptides are 14 aa in length, while it is 12, 14, and 16 for non-canonical peptides, contaminants, and variants, respectively.

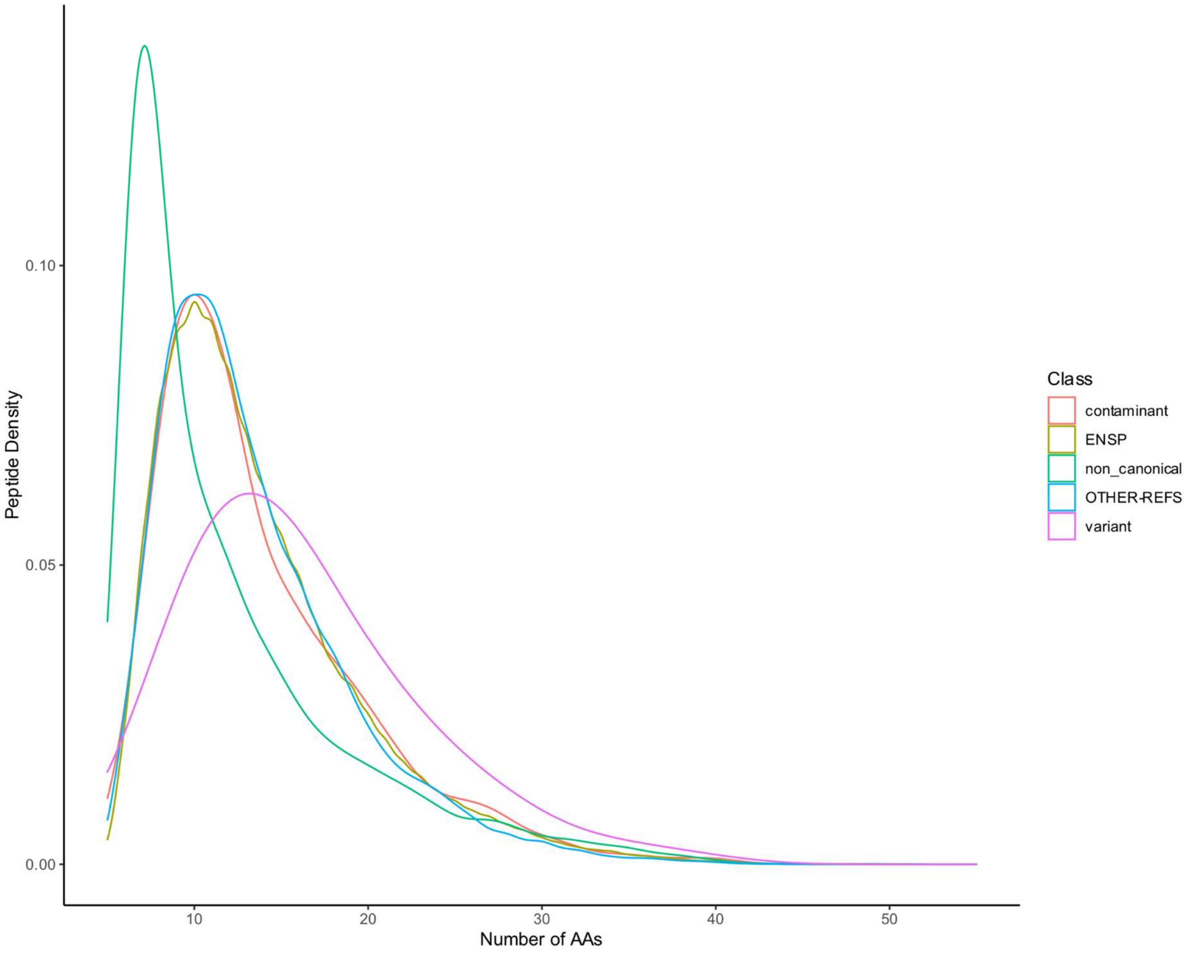

**Supplementary Note 4**: Distribution of novel peptide classes by cell lines

Distribution of number of novel peptides (noncanonical or variant) per cell line.

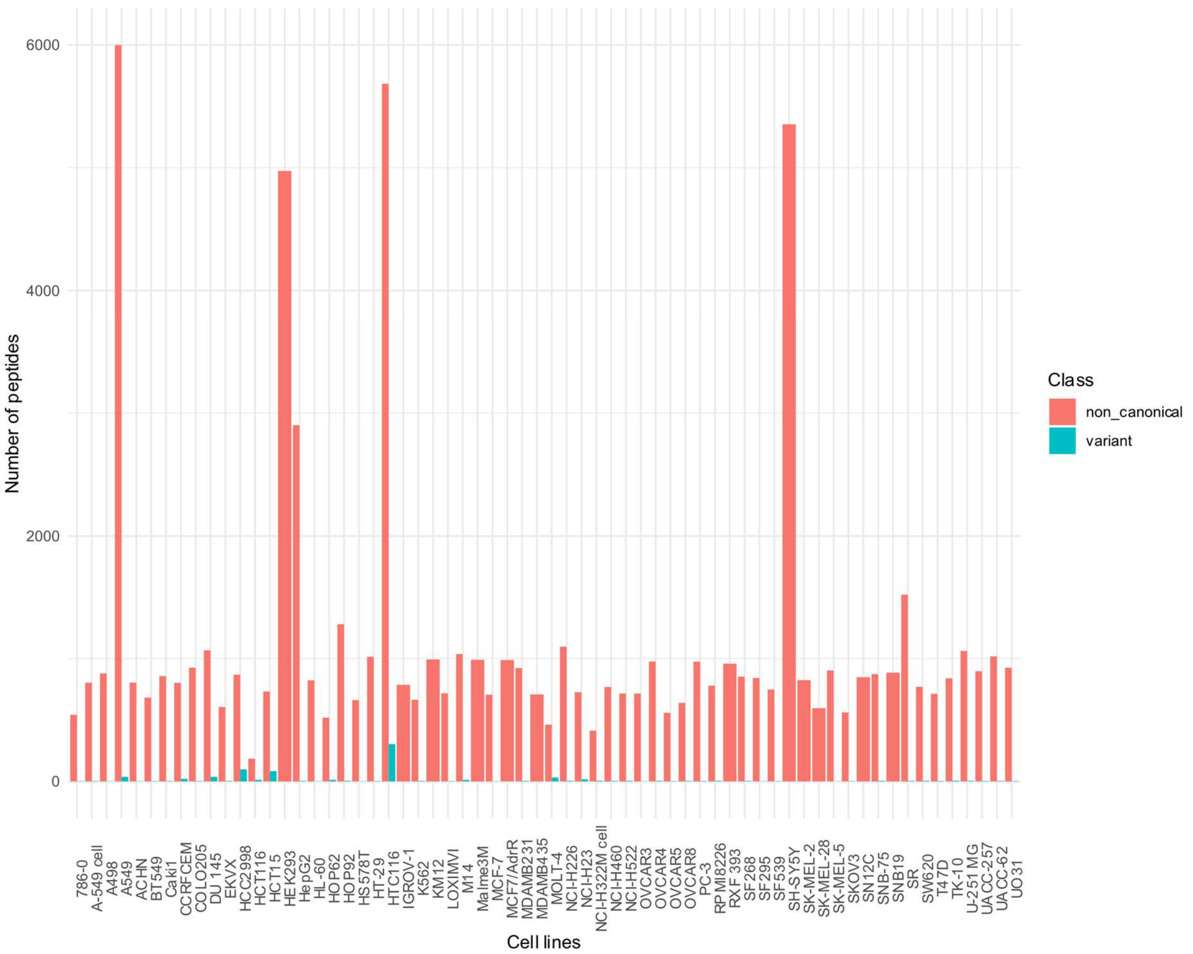
